# Cytokine and Growth Factor Response in a Rat Model of Radiation Induced Injury to the Submental Muscles

**DOI:** 10.1101/823922

**Authors:** Suzanne N. King, Zakariyya Al-Quran, Justin Hurley, Brian Wang, Neal Dunlap

## Abstract

Submental muscles (i.e. mylohyoid and geniohyoid) play a vital role during swallowing, protecting the airway from ingested material. To design therapies to reduce the functional deficits associated with radiation treatment relies in part on our understanding of the changes in the cytokine and growth factor response that can impact muscle function. The purpose of this study is to quantify changes in the inflammatory, pro-fibrotic, and pro-angiogenic factors following 48Gy of fractionated radiation to the mylohyoid muscle. We hypothesized that (1) irradiation will provoke increases in TGF-1β and MMP-2 mRNA in the mylohyoid muscle; and (2) muscles surrounding the target location (i.e. geniohyoid and digastric muscles) will exhibit similar alterations in their gene expression profiles. Rats were exposed to 6 fractions of 8Gy using a 6MeV electron beam on a clinical linear accelerator. The highest dose curve was focused at the mylohyoid muscle. After 2-and 4-weeks post-radiation, the mylohyoid, geniohyoid, and digastric muscles were harvested. Expression of TNF-α, IFNγ, IL-1β, IL-6, TGF-1β, VEGF, MMP-2, and MMP-9 mRNA was analyzed via PCR and/or RT-PCR. TGF-1β, MMP-2, and IL-6 expression was upregulated in the irradiated mylohyoid compared to nonirradiated controls. No notable changes in TNF-α, IFNγ, and IL-1β mRNA expression was observed in irradiated muscles. Differing expression profiles were found in the surrounding muscles post-radiation. Results demonstrated that irradiation provokes molecular signals involved in the regulation of the extracellular matrix, which could lead to fibrosis or atrophy in the swallowing muscle after radiation.

## Introduction

Radiation induced injuries are a major clinical concern after treatment for head and neck cancer, resulting in muscle weakness and impaired muscle mobility^1,2^. Previous clinical studies have proposed that these changes to the skeletal muscle after radiation are caused by prolonged inflammation that, in turn, leads to fibrosis and reduced mobility. This view comes from extrapolation of knowledge gleaned from the literature on radiation either to the oral cavity where destruction of the epithelial lining occurs or when high single doses are applied to the muscle at toxic levels. However, the molecular events underway in the irradiated skeletal muscle that are likely responsible for the development of swallowing mobility problems remain poorly understood. Evidence from other areas of the body suggest that radiation-induced injury to normal tissue is a dynamic and continuous process that is driven by the perturbation of multiple cells types (i.e. vascular, parenchymal, mesenchymal) within the tissue^3^. Cytokines and growth factors provide intercommunication between these participating cell types and changes in their expression can alter normal cell function. As such, further work is needed to identify key factors in the pro-inflammatory, pro-angiogenic and pro-fibrotic response within the irradiated swallowing muscles, which can negatively affect muscle regeneration.

The submental muscles of the floor of mouth, such as mylohyoid and geniohyoid muscles, are highly susceptible to radiation-associated impairments in swallowing function^4,5^. The mylohyoid facilitates elevation of the hyolaryngeal complex, while the geniohyoid provides anterior displacement of the complex. Together they prevent aspiration of food/liquid and aid in opening of the upper esophageal sphincter^6,7^. Injury to the mylohyoid muscle has been shown to cause kinematic changes in tongue displacement during licking and alters bolus transit during swallowing^8^. Recent animal studies have shown that radiation injury to other cranial muscles (i.e. genioglossus or hyoglossus) effects their contractile properties and increases their fatigability^9,10^. Molecular changes within the tissue may be contributing to these muscle injury induced mobility problems.

There is a paucity of research regarding the cellular and molecular responses following radiation injury to the skeletal muscles of the head and neck. Various cytokines and growth factors have been implicated in the pathogenesis of radiation injury including markers of inflammation (i.e. Interleukin [IL]-1β, IL-6, and tumor necrosis factor [TNF]-α), matrix remodeling (i.e. transforming growth factor [TGF]-β1) and vascular damage (vascular endothelial growth factor [VEGF]). In the skeletal muscle, increases in IL-1β, TNF-α, IL-6 and other inflammation-related molecules have been found within 4 to 6 hours of irradiation to abdominal muscles with a single dose of 10Gy^11^. In irradiated diaphragm muscles, elevation in oxidative stress has been observed 1-week after a single radiation dose of 10Gy, which can influence cell signaling and mediate repair. Fractionated radiation is thought to greatly enhance this cellular stress response, resulting in prolonged cytokine cascades compared to single doses^12^.

The matrix surrounding the muscle fiber provides structural support and protection; thus, any changes in its extracellular matrix (i.e. fibrosis or atrophy) may alter a muscles function. TGF-β1, VEGF and matrix metalloproteinases (MMPs) are actively involved in the turnover of extracellular matrix components following muscle injury. They also modulate endothelial cell function, promoting proliferation and angiogenesis needed for vascular repair. Of these cytokines, TGF-1β plays a pivotal role in normal tissue injury after radiation through its capacity to regulate numerous cell functions (e.g. growth, migration, matrix synthesis, and angiogenesis). In human samples, increases in TGF-β1 expression has been found in cervical strap muscles 4-weeks after radiation for oral or oropharyngeal cancers^13^. The overexpression of TGF-1β can lead to accumulation of collagen. This pro-fibrotic response is controlled by the synthesis of MMPs^14^. Two specific MMPs have been shown to be regulated by changes in contractile properties of the muscle, MMP-2 and MMP-9. Both of these MMPs degrade collagen type IV, which is a major component of the basement membrane of the skeletal muscle and facilitates the cellular arrangement of muscle fibers. Further work is warranted to determine the molecular changes undergone in the mylohyoid and surrounding muscles following radiation.

The objective of this study was to quantify changes in cytokine and growth factor expression following radiation to the mylohyoid muscle. We utilized a clinical linear accelerator to administer 48Gy of radiation given in 6 fractions of 8Gy to the rat, with the highest dose curve focused at the mylohyoid muscle. PCR and/or RT-PCR was performed with mRNA isolated from irradiated mylohyoid, geniohyoid, and digastric muscles at 2-and 4-weeks post treatment. We hypothesized that irradiation increases TGF-1β and MMP-2 mRNA expression within the mylohyoid. We anticipated that the inflammatory cytokine response would be minimal at the time points studied, indicating that there is no ongoing inflammation in the irradiated muscle. Additionally, we analyzed the gene expression patterns of the digastric and geniohyoid muscles, which surround the mylohyoid muscle of the rat, and were within the radiation field (albeit at lower doses). We hypothesized that muscles surrounding the target location will exhibit similar changes in their gene expression pattern.

## Methods

### Radiation Injury Model

Experiments involved nine adult male Sprague Dawley rats (~450-500kg). Six animals received radiation treatment as described below. Three animals were used as controls and underwent sham irradiation involving anesthetizing and immobilization on the platform for a similar length of time as the radiation group. All experimental protocols were approved by the Institutional Animal Care and Use Committee of the University of Louisville.

A clinically relevant radiation dose scheme was devised based on previous reports denoting that higher total dose volumes to the submental muscles are at greater risk for incidence of dysphagia^4,15^. Rodents were anesthetized with isoflurane (2-3%) via mask inhalation and placed in supine position on a customized platform to immobilize the animal. Rats underwent irradiation using a Clinac iX (Varian Medical Systems, Palo Alto, CA, USA) with 6 MeV electrons to limit the depth of maximum dose in the tissue to ~1.3-1.5 cm from the surface. To position the mylohyoid muscle at the depth of maximum dose, a 0.5 cm layer of tissue equivalent bolus was placed on the outer surface of skin above the submental space. A 6×6 cm electron applicator with a standard 4×4 cm insert was used to collimate the electron beam. A 0.3 cm lead shield was placed anterior to the bolus to further collimate the beam to the dimensions of the mylohyoid muscle (~11x 10mm). The distance from the radiation source to the surface of the bolus was 100 cm and the electrons were delivered at a dose rate of 1000 monitor units/minute. The number of monitor units needed to deliver the prescribed dose was calculated using an output factor for the 4×4 cm insert, which was measured by a clinical physicist. Boundaries and depth were determined based on gross anatomical dissections of the rat. Submental muscles were exposed to a total of 48Gy of radiation given in 6 fractions of 8Gy across a two week period. Oral intake and body weight were monitored to determine changes in overall well-being. One of the control animals received a 3-day course of antibiotics and nutritive supplements (Dietgel 76A) due to a torn toe nail.

### RNA Isolation

The mylohyoid, geniohyoid and digastric muscles were excised, placed in RNAlater and stored at −80□C for gene analysis studies. Total RNA was extracted from the muscles using SV Total RNA Isolation System (Promega Corp) according to the manufacturer’s instructions. First strand cDNA was synthesized from 1.5 μg of total RNA using an Omniscript Reverse Transcription Kit (Qiagen) and random primers (Integrated DNA Technologies, Coralville, IA).

### Polymerase Chain Reaction

To determine the gene expression profile of the mylohyoid muscle 2-and 4-weeks after irradiation, we measured TNF-α, IFNγ, IL-1, TGF-1β (all from^16^), IL-6, VEGF^17^, MMP-2^18^, and glyceraldehyde 3-phosphate dehydrogenase (GAPDH)^19^ mRNA. GoTaq Hot Start Polymerase (Promega Corporation, Madison, WI) was used with the PTC-300 Peltier Thermal Cycler according to the manufacturer’s protocol. Primer sequences, gene bank access numbers, and expected PCR product sizes are listed in Table 1. Specificity of each primer pair was confirmed by showing a single peak and appropriate sized DNA band for each gene product. Amplification was optimized for each primer and carried out for 30 cycles as previously described^20^. Reaction yields were determined by electrophoresis on 2% agarose gel using 12μl of each PCR product and visualized by ethidium bromide staining. Spleen samples were used as positive controls and included in each experiment. To establish a standard curve for RT-PCR (below), DNA fragments from spleen samples of amplified products were excised from the gel and purified using QIAquick Gel Extraction Kit (Qiagen) following manufacturers protocol.

**Table 1.**
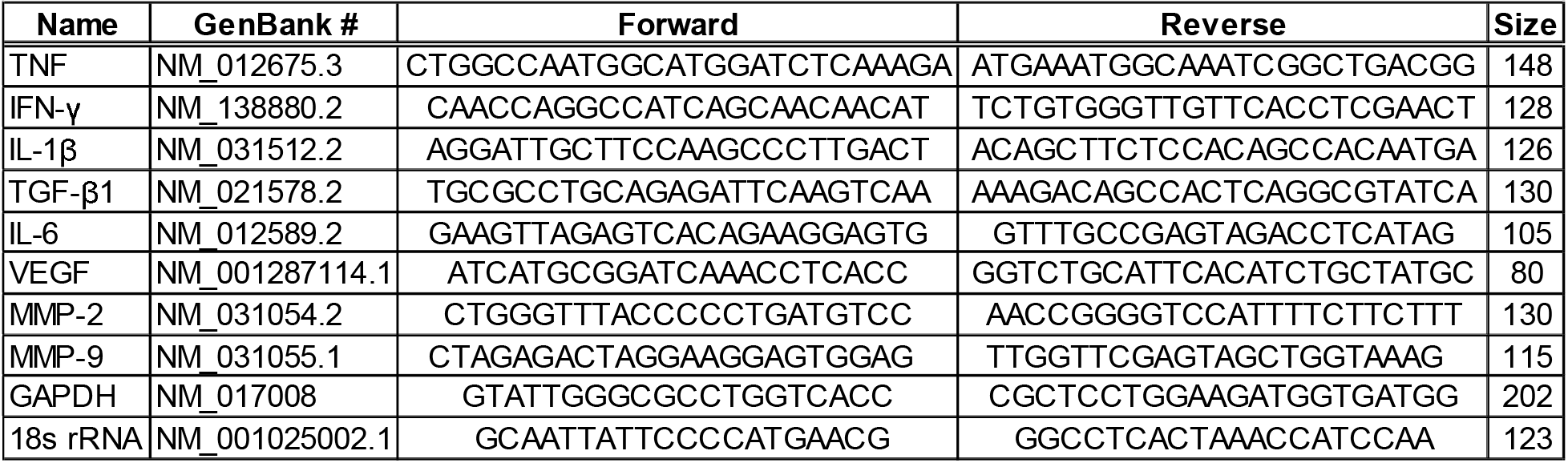
Primer Sequence and Products of Polymerase chain reaction (PCR)

### Real-time Polymerase Chain Reaction

To measure quantitative changes in genes expression in the mylohyoid, geniohyoid, and digastric muscles after irradiation, mRNA was measured utilizing relative quantification standard curve approach. Genes found to be expressed with PCR were further quantified using RT-PCR, including: IL-6, TGF-1β, VEGF, MMP-2, and MMP-9. SYBR Select Master Mix (Life Technologies) with the Applied Biosystem 7500 Fast System was used to quantify samples. GAPDH, 18s rRNA, and Beta 2 Microglobulin were tested for use as housekeeping genes. GAPDH was found to be the least effected by the experimental conditions. Negative (H2O) and positive (spleen) samples were included with each run. Eight-point standard curves were derived for each target to quantify gene expression and concentrations were normalized to the housekeeping gene.

### Statistical Analysis

A Welch’s one-way analysis of variance (ANOVA) was carried out to test the differences in time with each gene. Data was normally distributed, but violated the assumption of homogeneity of variances as assessed by Levene’s Test of Variance. If the mean differed significantly by p-value <0.05, a Games-Howell post hoc test was performed to compare groups. p<0.05 was considered statistically significant. All experiments were performed in triplicate. Data represents the mean ± standard deviation (SD).

## Results

### No noticeable changes in inflammatory genes in irradiated mylohyoid muscle

To characterize the gene response associated with radiation, cDNA from the mylohyoid at 2-and 4-weeks post was amplified for the target genes and analyzed using gel electrophoresis. Irradiated mylohyoid muscles demonstrated increased expression of IL-6, TGF-1β, VEGF, and MMP-2. No noticeable product was present for TNF-α, IFNγ, and IL-1β after radiation (Figure 1). All targeted genes were absent in normal mylohyoid tissue.

**Figure 1.**
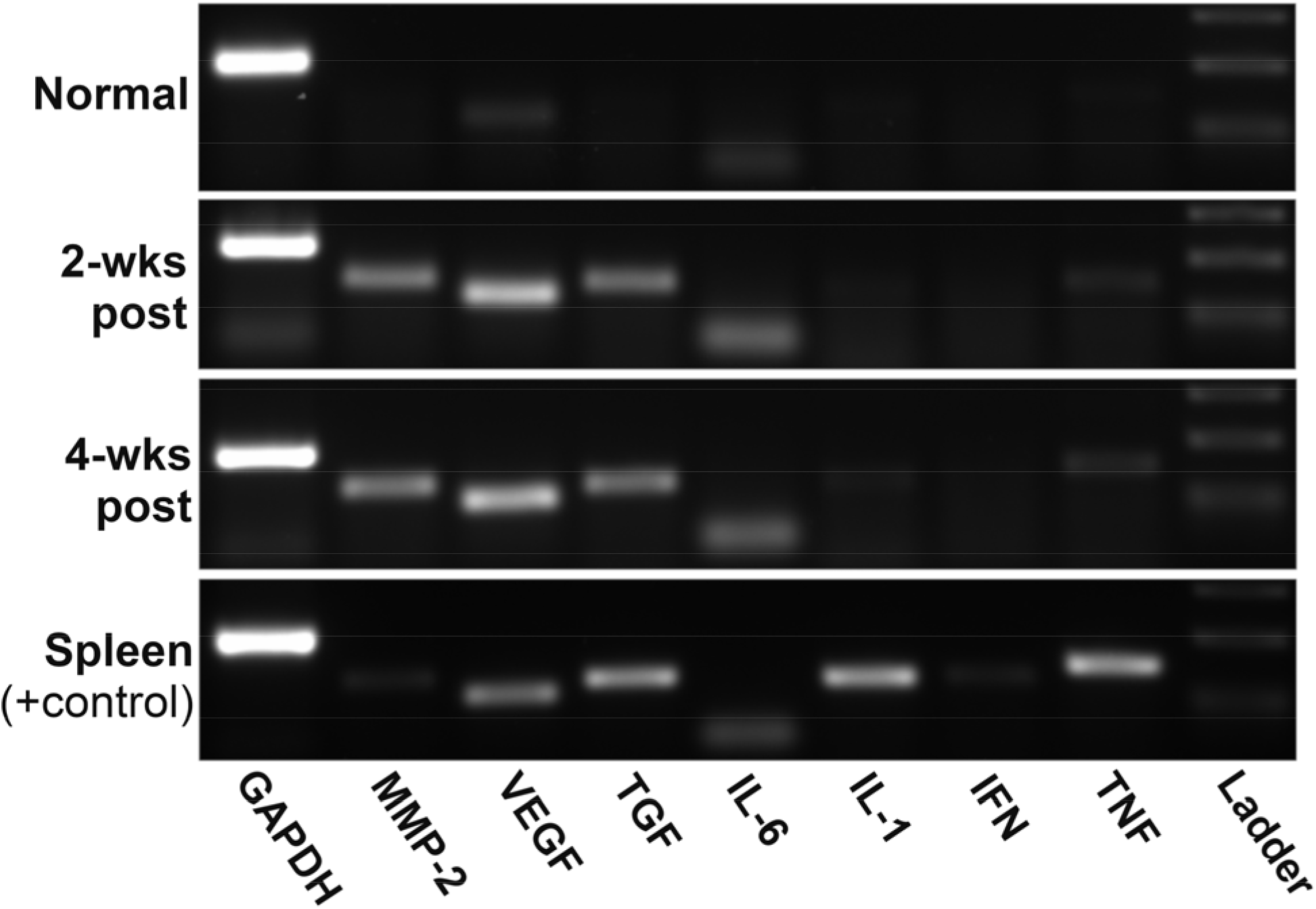
Gene expression of MMP-2, VEGF, TGF-β1, IL-6, IL-1, IFN, TNF, and GAPDH in mylohyoid muscles 2-and 4-wks post-radiation compared to normal controls. Polymerase chain reactions showed that the irradiated mylohyoid expressed MMP-2, VEGF, IL-6 and TGF-β1 mRNA compared to normal controls. Amplification of GAPDH was used as an internal control. Appropriately sized DNA band for each gene product was determined by 100 bp ladder. Spleen was used as a positive control.

### Differences in pro-fibrotic genes after irradiation

To quantify the differences in the pattern of expression, we analyzed cDNA from the mylohyoid, geniohyoid, and digastric muscle with RT-PCR. Only genes found to be expressed in mylohyoid with PCR were further analyzed with RT-PCR. Concentrations for each gene were determined by the relative standard curves and normalized to the housekeeping gene by calculating the ratio of the target gene to the housekeeping gene (ΔC_T_ Method). Data is presented in Figures 2–4 as the relative quantification.

**Figure 2.**
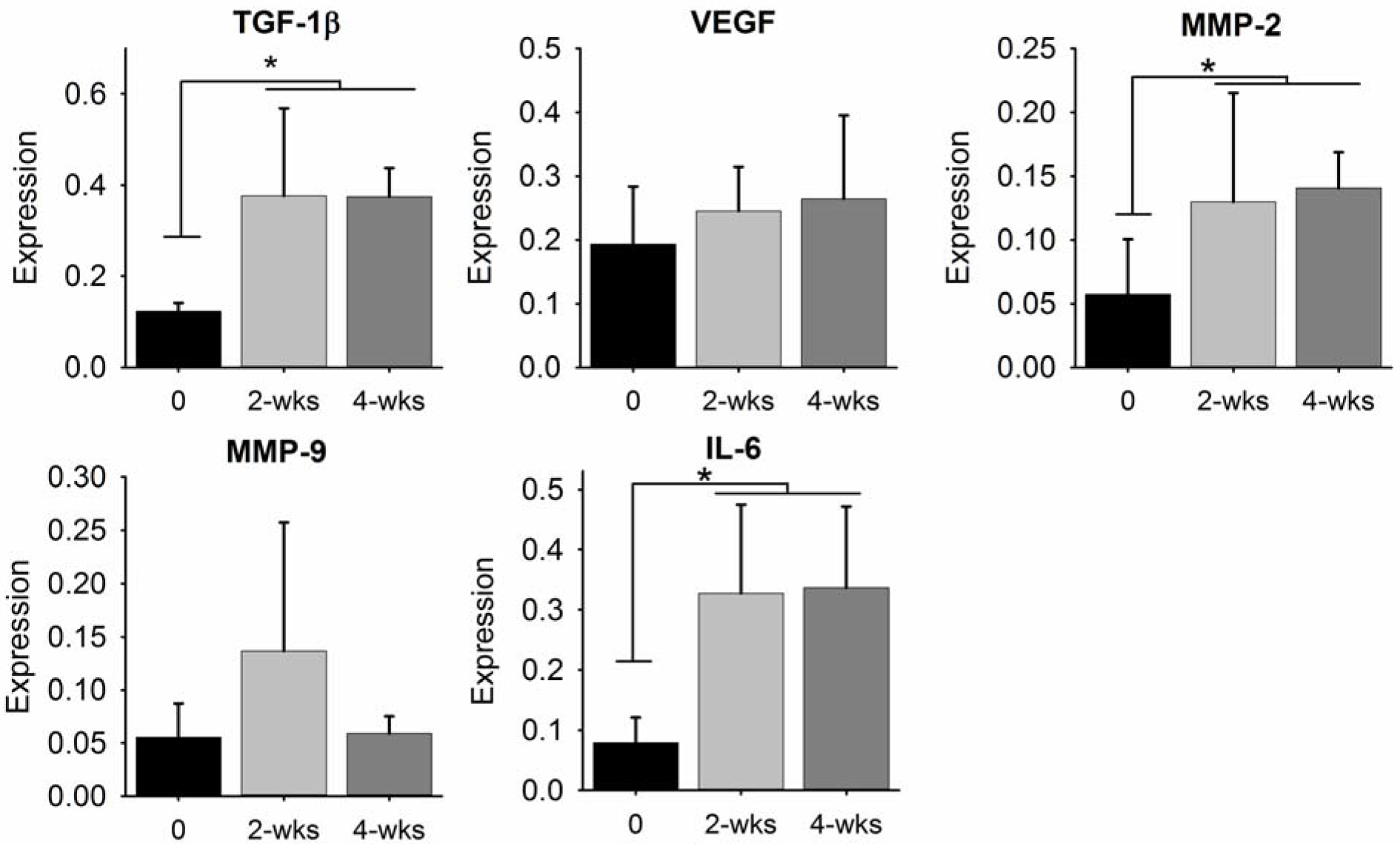
Comparisons of normal and irradiated mylohyoid muscle gene expression. Muscle was harvested and RNA was subsequently extracted, converted into cDNA and analyzed for TGF-β1, VEGF, MMP-2, MMP-9, and IL-6 gene expression using RT-PCR. Standard curves were derived to quantify gene expression and concentrations were normalized to GAPDH housekeeping gene. * represents a statistical significance of p< 0.05.

**Figure 3.**
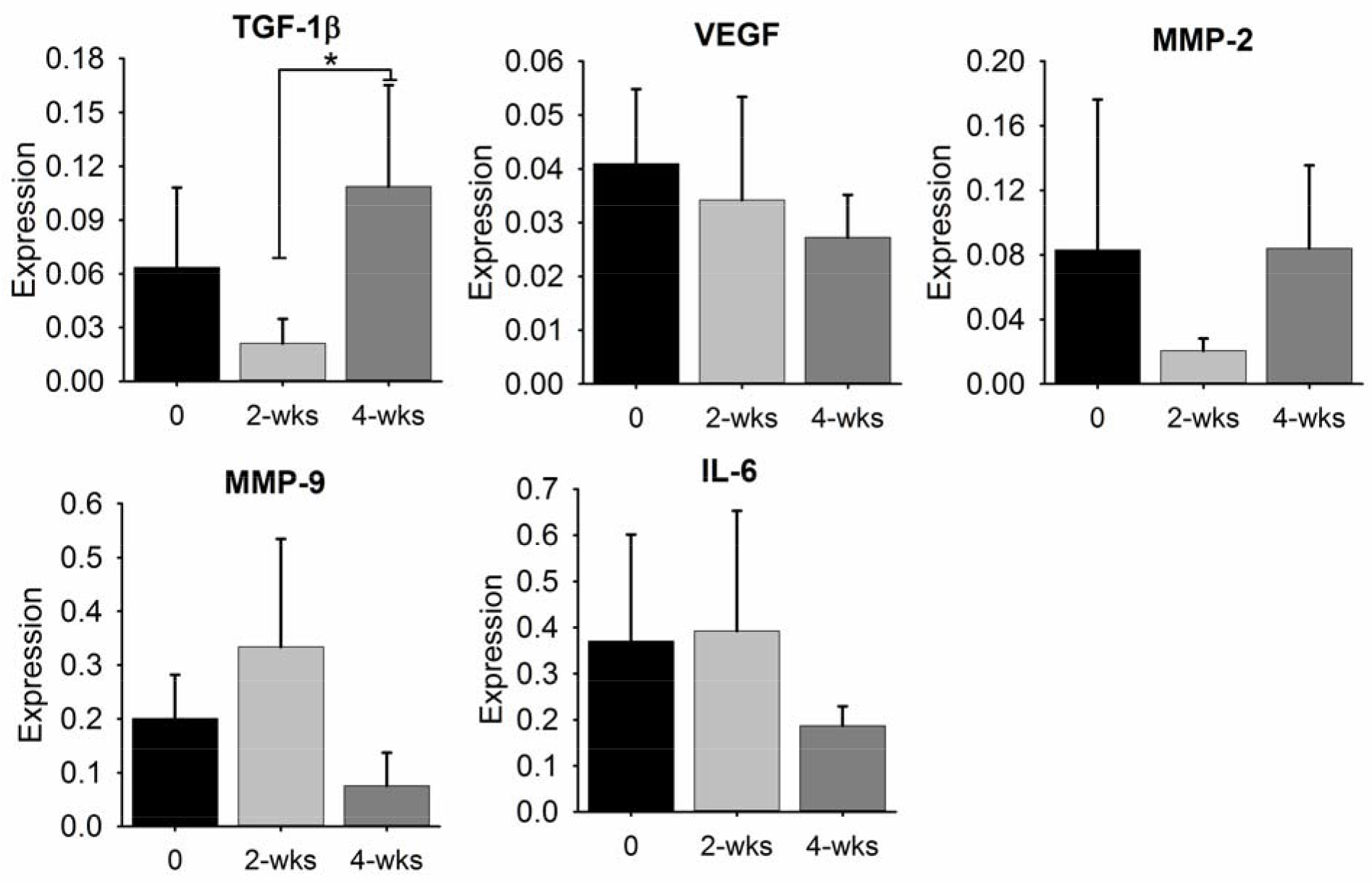
Comparisons of normal and irradiated geniohyoid muscle gene expression. Muscle was harvested and RNA was subsequently extracted, converted into cDNA and analyzed for TGF-β1, VEGF, MMP-2, MMP-9, and IL-6 gene expression using RT-PCR. Standard curves were derived to quantify gene expression and concentrations were normalized to GAPDH housekeeping gene. * represents a statistical significance of p< 0.05.

**Figure 4.**
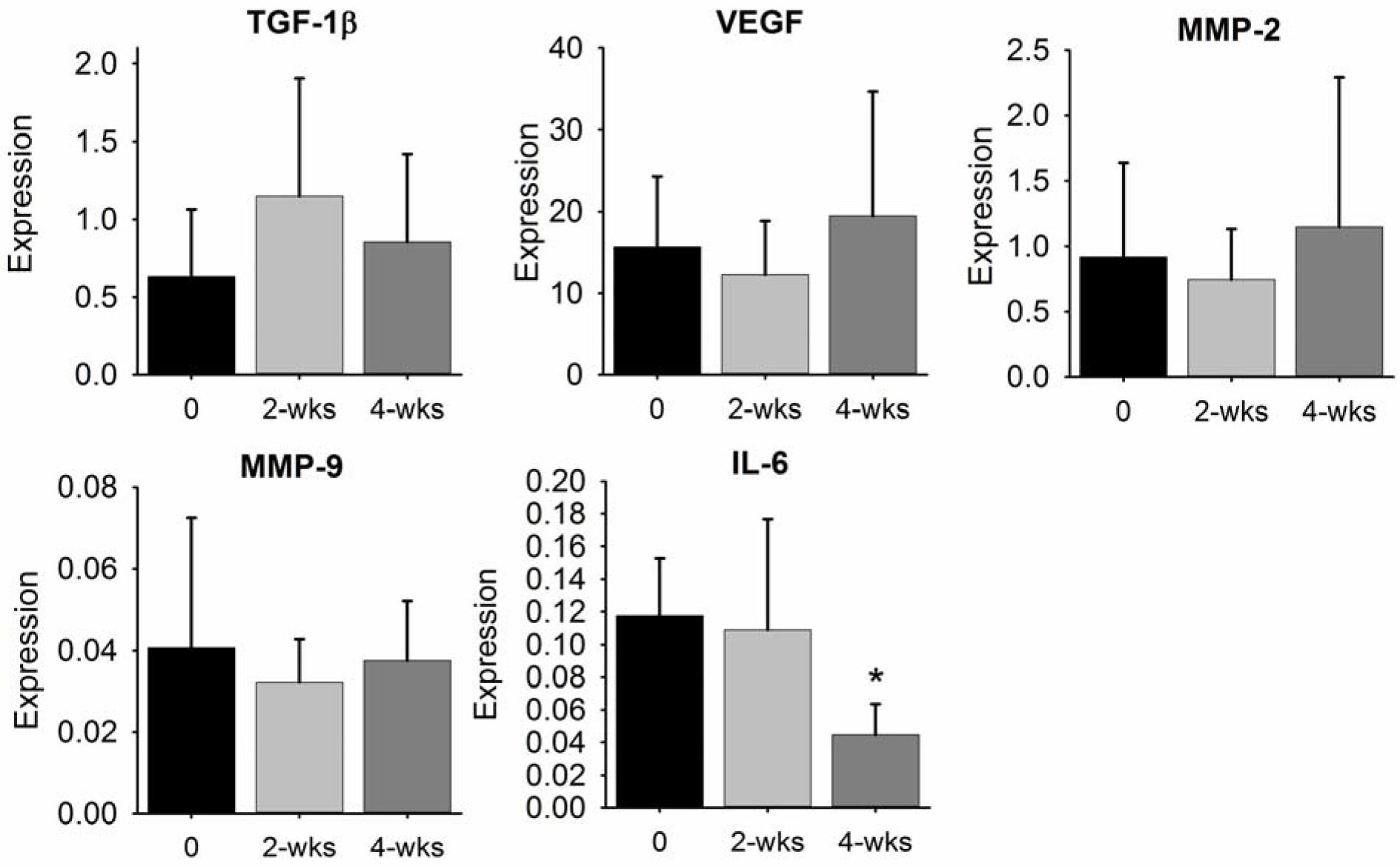
Comparisons of normal and irradiated geniohyoid muscle gene expression. Muscle was harvested and RNA was subsequently extracted, converted into cDNA and analyzed for TGF-β1, VEGF, MMP-2, MMP-9, and IL-6 gene expression using RT-PCR. Standard curves were derived to quantify gene expression and concentrations were normalized to GAPDH housekeeping gene. * represents a statistical significance of p< 0.05.

In the mylohyoid, radiation significantly affected the expression of TGF-1β (F[2, 7.156] = 40.838, p<0.0001), MMP-2 (F[2,9.105] = 8.201, p<0.009), and IL-6 (F[2,7.741] = 17.130, p<0.001). Significant increases in TGF-1β mRNA expression were found 2−(p<0.02) and 4-weeks post (p<0.001) compared to non-irradiated mylohyoid. MMP-2 was significantly increased 2-and 4-weeks after irradiation compare to non-irradiated controls (p< 0.05; p<0.02). IL-6 was significantly increased both at 2-and 4-weeks post-irradiation compared to controls (p<0.02; p<0.01). No significant differences were found in VEGF (p>0.4) or MMP-9 expression (p>0.3).

In the geniohyoid muscle, main affects in the expression of TGF-1β (F[2, 7.556] = 8.032, p<0.013), MMP-2 (F[2,6.848] = 5.251, p<0.041), and MMP-9(F[2, 9.00] = 7.155, p<0.014) were found after irradiation. TGF-1β significantly increased 4-weeks post radiation compared to 2-weeks post (p<0.03). MMP-9 was significantly higher in normal controls compared to 4-weeks post radiation (p<0.04). No significant post hoc differences were found with MMP-2, VEGF, or IL-6 in the geniohyoid muscle.

In the digastric muscles, significant changes in IL-6 (F[2, 8.384] = 10.425, p<0.005) were observed following radiation. IL-6 significantly decreased 4-weeks after radiation compared to 2-weeks post (p<0.02) and controls (p<0.01). No significant differences were found in TGF-1β, MMP-9, MMP-2, or VEGF in irradiated digastric muscles.

## Discussion

There is much confusion in the literature regarding the molecular response to radiation injury in the submental muscles. The majority of the research on radiation to the head and neck focus on deficits in the oral cavity where the epithelial lining is destroyed resulting in ulcerations, mucositis and other significant dose limiting effects. The submental muscles lye deep to the submandibular space and are covered superficially by the dermis and platysma muscles and thus, not likely directly affected by a mucosal injury. In an effort to clarify the cytokine and growth factor response induced by radiation treatment to the submental muscles during the subacute period, we examined the mylohyoid, geniohyoid, and digastric muscles, which are vital swallow-related muscles involved in the movement of the hyolaryngeal complex. Following 48Gy of radiation to the mylohyoid given at 8Gy for 6 fractions, we found a predominate pro-fibrotic response, suggesting a potential role for these cytokines in the induction of radiation muscle injury. Our data revealed an increase in TGF-β1 and MMP-2 was independent of an ongoing inflammatory response in the mylohyoid muscle.

There is overwhelming evidence for involvement of TGF-β1 in response to radiation irrespective of the tissue location^21^. Our findings confirmed its upregulation in the irradiated mylohyoid and geniohyoid after treatment. TGF-β1 signaling regulates diverse biological activities through SMAD-dependent pathways, as well as other transcription pathways i.e. MAPK and NF-kB. Overexpression of TGF-1β has been linked to fibrosis and atrophy post muscle injury. The onset of oxidative stress and pro-inflammatory stimuli rapidly after radiation exposure likely act as an initial trigger for activation of TGF-1β^22^. In the rat, expression of TGF-β1 has been shown to increase after radiation to liver and lung in a dose-dependent manner and its expression correlates with the upregulation of collagen^23,24^. However, previous reports in other cranial muscles have found no changes in collagen concentration (at the protein level) after irradiation, even though changes in motor function were observed. Russel et al, irradiated the genioglossus muscle of the rat (11Gy × 2) and found no observable changes in collagen content after 6-months post treatment^9^. Benedict et al treated the tongue base with chemoradiation (7Gy × 5) and also, found no changes in collagen within the hyoglossus muscle 2-weeks and 5-months post^10^. This suggests that other molecular changes in the muscle are likely hindering movement of the irradiated cranial muscles. TGF-β1 also acts as a potent inhibitor of muscle regeneration and promotes muscle weakness/atrophy. Reports have shown that increases in TGF-β1 (via local delivery) into the normal hindlimb muscle of a mouse leads to muscle atrophy including a loss of muscle mass and decreases in muscle fiber size^25^. Other studies have also shown that overexpression of TGF-β1 at early time points post muscle damage, suppresses myogenic response (i.e. MyoD and myogenin) through SMAD3 signaling pathways, leading to failure of the injured muscle to properly regenerate^26^. Because TGF can have a multitude of functions depending on the injury, time period, and muscle it is effecting, additional studies are warranted to determine its contribution to the pathogenesis of radiation injury to the swallowing muscles.

Activation of MMP-2 and MMP-9 play a pivotal role in regeneration after inflammatory induced changes in skeletal muscle and exhibit differences in response to injury^27^. MMP-9 is typically found early after muscle injury, provoked by inflammatory cells i.e. macrophages in an effort to regulate matrix turnover. MMP-2 expression increases during normal muscle remodeling, facilitating the degradation of collagen types IV, VII and X^28,29^. In the current study, we found that radiation to the mylohyoid showed a distinct upregulation in MMP-2 mRNA expression, with no observable changes in MMP-9. Because MMP-2 facilitates the degradation of collagen within the muscle, upregulation in the irradiated mylohyoid could affect its structural integrity. Prior work in a rat model suggested that muscle atrophy may be attributed to an upregulation in MMP-2, based on increases in both collagen degradation and MMP-2 production following immobilization of the hindlimb^30^. Previous reports have also implicated that the proportion of slow and fast twitch fiber types effects the pattern of MMP-2 and MMP-9 expression in the muscle and therefore, attributes to the effectiveness of muscle regeneration after injury^31^. For instance, fast twitch muscles (i.e. extensor digitorum longus), which exhibit improved regeneration capacity, express higher MMP-2 activity and lower MMP-9 expression than slow twitch muscles (i.e. soleus) after injury. However, in smaller animals (i.e. rat) about 90% of the fibers in the submental muscles are fast contracting, type II myosin heavy chain (MHC) isoforms^32^. Thus, there are fewer slow contracting fibers in the rat submental muscles compared to larger animals and humans. In the normal rat mylohyoid muscle, fibers consist of primarily fast type MHC isoforms (i.e. type IIx), which is much higher than the ~50% detected in humans^33^. In the normal rat geniohyoid and anterior digastric muscles, there are predominately fast-contracting, anaerobic MHC type 2a and 2b fibers^32,34^. Therefore, it is unclear from our study if fiber type is regulating the MMP expression levels found. Of note, the enzymatic activity of MMP-2 is regulated at the level of synthesis by tissue inhibitors of metalloproteinases (TIMP). However, there is no previous evidence indicating that TIMP expression is regulated by radiation injury. Prior work in the lung observed no differences in TIMP expression following radiation^29^. Further work is needed to determine if increased expression of MMP-2 in the mylohyoid is associated with muscle atrophy.

Following radiation, we found that IL-6 mRNA was elevated in the mylohyoid, but no differences were observed in the other surrounding muscles. A decrease in oxygen availability, called hypoxia, has been shown to regulate IL-6 and VEGF levels after irradiation in lung and rectum. Chronic hypoxia in the skeletal muscle results in vascular damage and in turn, stimulates the upregulation of various cytokines by vascular endothelial cells^35^. However, the expression of VEGF in submental muscles differed from the results obtained in other areas of the body, which found this growth factor to increase after radiation^36,37^. In these studies they used high single doses of >25Gy to induce injury. For example, Liu et al observed max expression of VEGF 4-weeks after a single dose of 25Gy to the rectum of the mouse. It is possible that the hypoxic response to radiation is dose dependent. Further work is needed to determine if hypoxia is induced following irradiation with higher total radiation doses to the swallowing muscles.

There are limitations of this study that warrant further discussion. Because we analyzed homogenated muscle samples, we could not distinguish the cell type that was increasing the local production of the cytokines and growth factors. For instance, IL-6 could have been expressed by vascular endothelial cells and/or muscle fibers in response to irradiation. Future studies should isolate the mechanism and cellular source of the cytokine contributing to these pathologic changes. Secondly, the results presented at the 2-week time point contained significant individual variability, likely due to differences in the biological response to radiation. Our objective was to study the subacute changes associated with radiation-induced injury. In humans, this is thought to occur between 2-6 weeks post treatment. Further work at early time points are warranted to further identify factors that may influence the negative effects on muscle function after radiation. Lastly, our radiation injury model focused the highest radiation dose to the mylohyoid and thus, our results on the digastric and geniohyoid muscles reflect lower radiation doses as the dose curve rapidly drops off. This is likely why we observed differences in gene expression between the muscles tested. Interesting, although geniohyoid received a lower radiation dose, TGF-1β expression was still found to be upregulated at 4-weeks post-radiation, possibly indicating that the temporal response to these genes may have a dose threshold.

## Conclusion

In summary, results demonstrated that radiation provoked a unique cytokine and growth factor responses in the mylohyoid, geniohyoid, and digastric muscles. Irradiation not only affected the targeted muscle, but surrounding muscles in the radiation field were also modulated to a lesser degree. Of particular interest was that irradiation stimulated molecular events involved in the regulation of the extracellular matrix, which could lead to fibrosis or atrophy in the muscle.

